# Precision weighting of cortical unsigned prediction errors is mediated by dopamine and benefits learning

**DOI:** 10.1101/288936

**Authors:** J. Haarsma, P.C. Fletcher, H. Ziauddeen, T.J. Spencer, K.M.J. Diederen, G.K Murray

**Author notes:** joint last author/equal contributions. Correspondence to: Dr Diederen, Department of Psychosis studies, Institute of Psychiatry, Psychology and Neuroscience King’s College London, De Crespigny Park, London, SE5 8AF, United Kingdom **or** Dr Murray, Department of Psychiatry, University of Cambridge, Douglas House, Trumpington Road 18b, Cambridge, CB2 8AH, United Kingdom.

## Abstract

The predictive coding framework construes the brain as performing a specific form of hierarchical Bayesian inference. In this framework the precision of cortical unsigned prediction error (surprise) signals is proposed to play a key role in learning and decisionmaking, and to be controlled by dopamine. To test this hypothesis, we re-analysed an existing data-set from healthy individuals who received a dopamine agonist, antagonist or placebo and who performed an associative learning task under different levels of outcome precision. Computational reinforcement-learning modelling of behaviour provided support for precision-weighting of unsigned prediction errors. Functional MRI revealed coding of unsigned prediction errors relative to their precision in bilateral superior frontal gyri and dorsal anterior cingulate. Cortical precision-weighting was (i) perturbed by the dopamine antagonist sulpiride, and (ii) associated with task performance. These findings have important implications for understanding the role of dopamine in reinforcement learning and predictive coding in health and illness.

## Introduction

Recent decades have witnessed an increased interest in Bayesian theories of neural function, where neural signals representing prediction errors (expected – predicted outcome) play a key role (Rao & Ballard, 1999; den Ouden et al., 2008 & 2012; Bastos et al., 2012). These models, including predictive coding frameworks and the free-energy principle, conceptualize the brain as performing a specific form of hierarchical Bayesian inference, where each layer in the cortex aims to best predict its input, and updates its prediction by virtue of the prediction error, to achieve an accurate model of the world (Rao & Ballard, 1999; Bar, 2009; Friston, 2009; Mathys et al., 2011; Bastos et al., 2012; Clark, 2013 & 2015; Hohwy, 2013). A core idea included in these models is that prediction errors are scaled in a precision-weighted fashion, i.e. neural systems encoding predictions errors respond more strongly when new information is more reliable and hence more informative. Neural implementations of these theories have hypothesized the importance of the dopamine system in controlling the precision-weighted coding of prediction errors (Friston, 2009; Bastos et al., 2012; Adams et al., 2013). However, to the best of our knowledge, there is no direct evidence to support the hypothesis of a dopamine-mediated precision-weighted prediction error signalling in the cortex.

As with Bayesian theories of neural function, reinforcement-learning models also highlight the importance of the minimization of prediction error (Sutton & Barto, 1998). However, in this context a distinction is usually made between signed and unsigned prediction errors, which play different roles (Roesch et al., 2012). The signed prediction error signals the extent to which an outcome is better or worse than expected and changes the value of a stimulus accordingly (Rescorla & Wagner et al., 1972; Sutton, 1988). The unsigned prediction error signals the degree to which an outcome is unexpected, independent of its sign, and thereby controls the rate of learning (Pearce & Hall 1980; Hall & Pearce et al., 1982). In Bayesian reinforcement learning models unsigned prediction errors increase (or decrease) the uncertainty regarding prediction estimates and thereby increase learning in a fashion similar to reinforcement learning (Courville et al., 2006). Critically, influential Bayesian models also posit the presence of a cortical precision-weighted unsigned prediction error signal, thought to be mediated by dopamine (Friston, 2009; Bastos et al., 2012; Adams et al., 2013).

Where in the brain can we expect to find evidence of a precision weighted unsigned prediction error signal that is mediate by dopamine? During reinforcement learning, neural signals representing signed reward prediction errors and motivational salience prediction errors can be found in various locations (Roesch et al., 2012). For example, signed reward prediction errors are coded in a variety of brain regions, most notably the dopaminergic midbrain and ventral striatum (Schultz et al., 1997; O’Doherty et al., 2003 & 2004; Pessiglione et al., 2006; D’Ardenne et al., 2008; Diederen et al., 2016 & 2017, Tian et al., 2016). In contrast, neurophysiological evidence in experimental animals suggests that unsigned prediction errors, which are the main focus of interest here, are coded in the cortex, including the dorsal anterior cingulate cortex (dACC) (Hayden et al., 2011), as well as subcortically, in the basolateral amygdala (Li et al., 2011); both of these structures are innervated by dopamine (Esber et al., 2012; Paus et al., 2001). Evidence from human fMRI learning studies also points to prefrontal representations of unsigned prediction error, in medial, superior and lateral aspects of the prefrontal cortex, including the dACC (Fletcher et al 2001, Turner et al 2004, Ide et al., 2013; Metereau et al., 2013; Fouragnan et al., 2017, Fouragnan et al 2018).

Given the findings of unsigned prediction error signals in the frontal cortex, and given that the dACC receives dense dopaminergic projections (Lewis et al., 1987; Berger et al., 1991), we hypothesize the following. Firstly, we expect that unsigned prediction errors are coded in the frontal cortex. Secondly, we expect that the unsigned prediction error signal is coded relative to the precision of environmental outcomes. Third, we expect the aforementioned process to be mediated by dopamine, especially in the dACC. We test these hypotheses with novel analyses on an existing dataset (Diederen et al 2017). In this study participants were randomised to receive either placebo, sulpiride (a D2 receptor antagonist), or bromocriptine, (a potent dopamine receptor agonist, which also has effects at some classes of 5-HT receptors (Knverno et al., 2008; Newman-Tancredi et al., 2002)). Participants undertook a cognitive task where they were presented with rewards/ monetary outcomes on each trial that were drawn from distributions with varying degrees of uncertainty. The participants were instructed to best predict drawn rewards, with the most accurate prediction corresponding to the mean of the reward distribution. Crucially, we manipulated the reliability of reward information by including distributions with varying standard deviations. Regression analyses clearly indicated a role for unsigned prediction errors in learning. Further computational modelling confirmed that behaviour was best explained by a model that included precision-weighted signed and unsigned prediction errors that drove value updating and learning rates respectively. Functional MRI (fMRI) in conjunction with a pharmacological manipulation allowed us to explore whether dopamine perturbs the precision-weighting of prediction errors in the brain, and whether this perturbation was associated with diminished task performance. Our previous analysis of this dataset focussed exclusively on analysis of signed prediction errors and was largely focussed on subcortical region of interest analyses (Diederen et al 2017). Specifically, these analyses showed precision-weighting of signed prediction error signals in the dopaminergic midbrain and striatum, the extent of which was modulated by D2 receptor antagonism. However, in the current paper, we are concerned primarily with cortical representations of unsigned prediction error, and their dopaminergic modulation. In a novel analysis, we find precision-weighting of unsigned prediction error signals in the superior frontal cortex and dACC, which are modulated by the dopamine D2 receptor antagonist sulpiride. The findings provide evidence in favour of recent computational models that hypothesize a role for dopamine in modulating precision-weighting of cortical prediction error signals (Bastos et al., 2012; Adams et al., 2013).

## Results

### Environmental precision and D2 antagonism modulate performance

First, we explored whether performance increases when environmental precision is high (i.e. when the standard deviation of rewarding outcomes is low). We measured performance by investigating how close participants predictions were to the actual mean of the distribution in the final trials of each task block (i.e., the average difference between the mean of the reward distribution and the final three predictions the participants made). We used a two-factor mixed model ANOVA with medication group as the between-subjects variable and SD conditions as the within-subjects variable, using a linear contrast across SD for the main effect of SD and interaction. We found that performance increased when the standard deviation decreased (F{1,56}=11.3, *p*=.001). We found no interaction between SD and group (F{2,56}=1.3, *p*=.27). However, there was a trend-level effect of group on average final performance (F{2,56}=2.5, *p*=.094). Post-hoc tests indicate reduced overall performance in the sulpiride group compared to placebo (F{1,38}=5.10, *p*=.030), but no significant interaction with SD (F{1,38}=.18, *p*=.68). Comparing the bromocriptine group to placebo there was no significant overall difference (F{1,37}=2.01, *p*=.17) and no interaction (F{1,37}=1.70, p=.20).

Since dopamine plays an important role in motivation, we explored whether the dopaminergic manipulation influenced measures of motivation, such as reaction times (Crespi, 1942; Niv, 2007), how far the participants scrolled the mouse in order to come to state their prediction (Fig. 1A; please see methods section for more detail), and the number of missed trials (Fig. 1D). Using one-way ANOVAs, we found that only reaction times were significantly different across medication groups, with the sulpiride group being faster than placebo (RT: F{2,55}=4.21, *p=*.019; Misses: F{2,55}=0.33, *p*=.72; Scrolling distance: F{2,55}=0.56, *p*=.57; Fig. 1C). Differences in RT remained significant when controlling for scrolling distance: F{2,55}=3.32, *p*=.044. More rapid reactions in the sulpiride condition makes it unlikely that any performance deficit secondary to sulpiride could be driven by motivational impairments.

**Figure 1:**
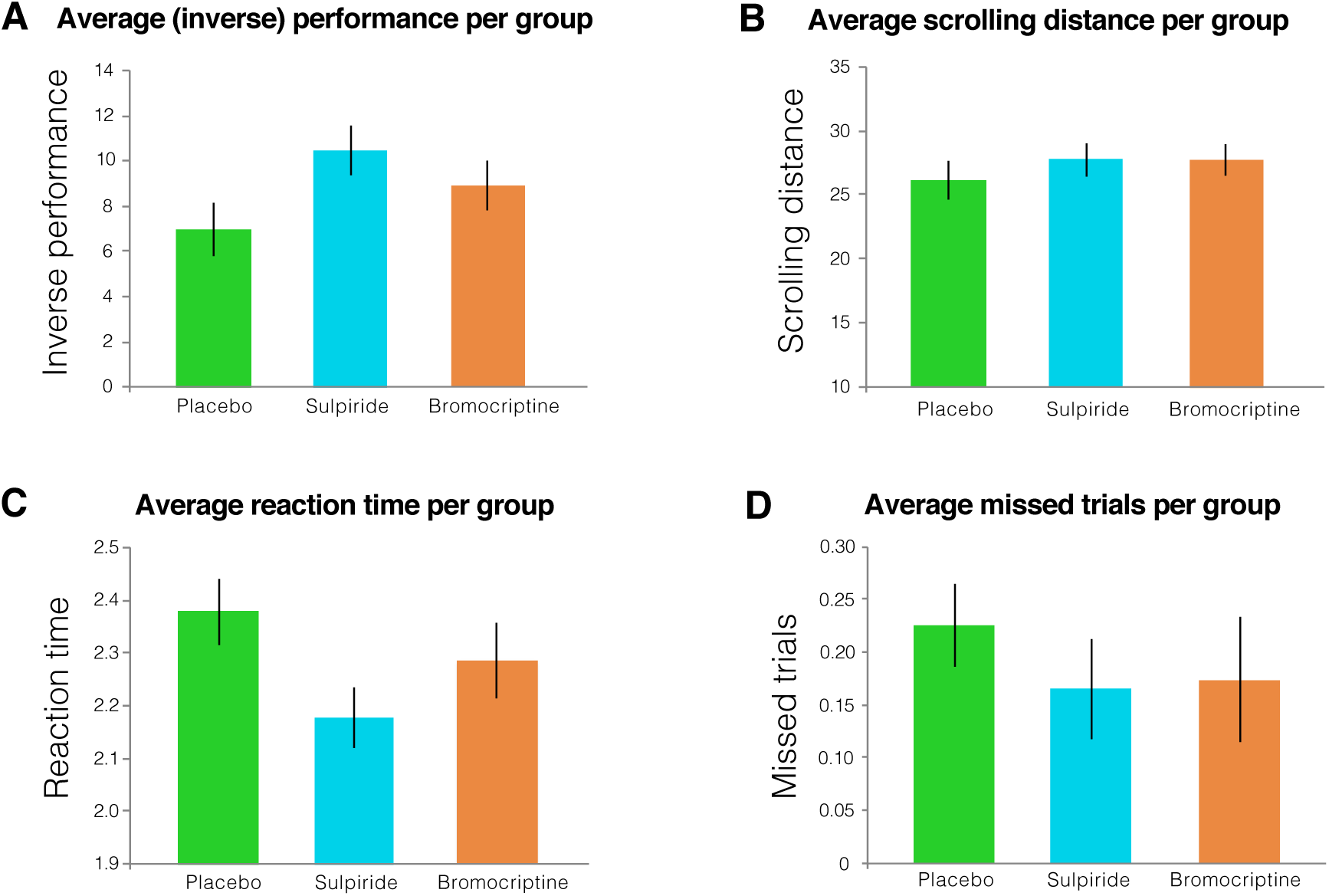
A. Average performance, B. scrolling distance, C. reaction time and D. missed trials per group. Error bars represent standard error of the mean. Participant performed worse after receiving sulpiride, and they responded significantly faster.

### Unsigned and signed prediction errors contribute to belief updating

In order to explore whether signed and unsigned prediction errors contribute to belief updating, we fitted a simple regression model to the behavioural data, where belief updating was the dependent variable and signed and unsigned prediction errors the independent variables. Belief updating here is defined as the change in predicted reward from t_n_ to t_n+1_. We expected a main effect of signed prediction error, but not unsigned prediction as the former informs the participants about the direction of belief updating, whereas the latter does not. However, as reinforcement learning models like the Pearce-Hall model suggest that unsigned prediction errors increase the amount of attention devoted to a stimulus, we expect there to be an interaction between unsigned and signed prediction errors. As expected we indeed found an effect of signed prediction errors (F{1,10763}=246.95, *p*<.0001), but not unsigned prediction errors (F{1,10763}=.95, *p*=.33). Most importantly we found an interaction effect between signed and unsigned prediction errors (F{1,10763}=51.7, *p*<.0001), showing that the influence of signed prediction error on belief updating is a function of both the signed and unsigned prediction error. We furthermore tested whether SD condition had an interaction with signed prediction error on belief updating. Although there was no main-effect of SD (F{2,10761}=.25, *p*=.78) there was indeed a significant interaction (F{2,10761}=25.8, *p*<.0001). In order to visualise the interaction between signed and unsigned prediction error we binned signed prediction errors and plotted belief updating on the y-axis (Fig. 2A). The significant interaction predicts a logit-function, amplifying the contribution of signed prediction error on belief updating when the unsigned prediction errors are highest (i.e. when signed prediction errors are highly positive or highly negative). We thus conclude unsigned prediction error contribute to learning in addition to signed prediction errors. We furthermore visualised the effect of SD, where we see that signed prediction errors have a stronger effect on learning in the SD5 condition compared to SD15 (Fig. 2B). The significant interaction predicts a stronger effect of signed prediction error on belief updating in the SD5 condition compared to the SD15 condition. We thus conclude signed prediction errors contribute more to learning when reward information is reliable.

**Figure 2:**
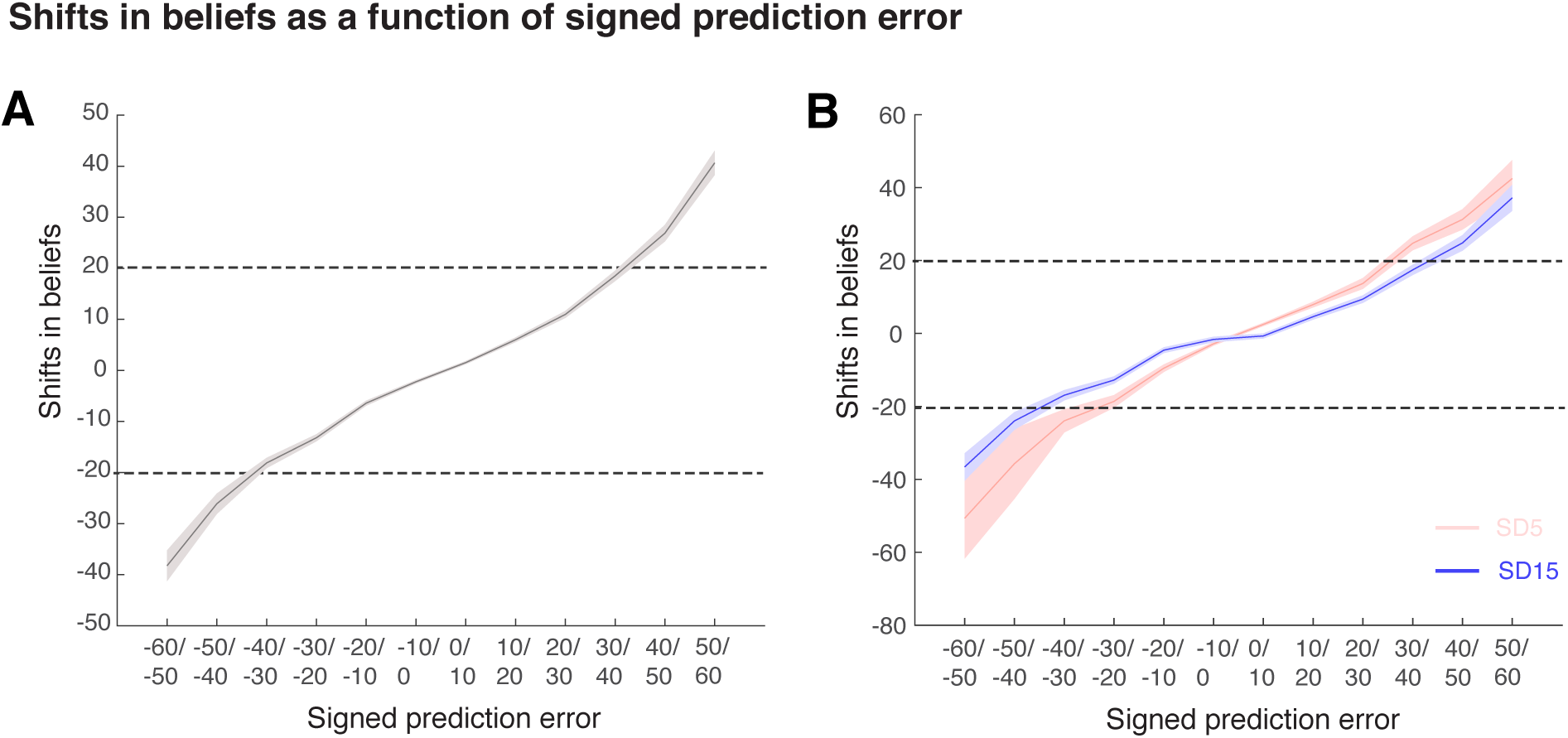
Here we show the relationship between signed prediction errors (x-axis) and belief-updating (y-axis) collapsed for SD5 SD10 & SD15 (A) and separate for SD5 & SD15 (B) (SD10 is not plotted for clarity purposes). The shaded area is one standard error of the mean. In fig2A the interaction effect between unsigned prediction error and signed prediction is shown (above the top and below the bottom dotted line unsigned prediction errors are strongest, amplifying the effect of the signed prediction error). In figure B the effect of SD is shown, revealing a stronger effect of signed prediction error in the SD5 condition compared to SD15.

### Reinforcement Learning modelling of the behavioural data supports a precision-weighted unsigned and signed prediction error

We fitted different reinforcement learning models to the participants’ trial-wise predictions to investigate whether participants were likely to make us of precision weighted unsigned and signed prediction errors. These models included a simple Rescorla-Wagner (RW, Rescorla & Wagner, 1972) model without precision weighting of prediction errors, as well as a precision weighted RW model. In addition, as previous work using similar tasks showed that learning rates tend to decrease as trials progress as a function of unsigned prediction error size in uncertain environments we included four Pearce-Hall (PH, Pearce & Hall, 1980) models that allow the learning rate to decrease as trials progress: 1) without precision weighting of prediction errors, 2) precision weighting of signed and unsigned prediction errors 3) estimated precision weighting of unsigned and signed prediction errors 4) separate estimation of precision weighting of unsigned and signed prediction errors. The modelling showed that in all three medication groups the Pearce-Hall model with separate estimated precision weighted signed and unsigned prediction errors performed the best in explaining the behavioural data (model comparison using AIC to penalize models for the number of free parameters; see table 2 for an overview of the model (free) parameters and models fits and comparisons). These results indicate that both unsigned and signed prediction errors are precision-weighted to facilitate efficient learning when the environment is uncertain/ variable.

### Unsigned prediction errors are coded in the SFC/dACC

Our previously published analysis of these neuroimaging data focussed exclusively on signed prediction errors (Diederen et al., 2017), and was largely confined to subcortical region of interest analysis. To investigate the coding of unsigned prediction errors in the brain, here we tested for a main effect on the unsigned prediction errors parametric modulators (regressors whose magnitude is proportional to the degree of unsigned prediction error). Note that the task design allows us to measure the degree of prediction error directly by taking the difference between predicted and received reward, and therefore do not need to be inferred via the modelling procedure. This has the added benefit that fMRI group effects cannot be driven by differences in model fits between groups.

We conducted analyses across the whole brain at p_FWE_ <.01. Unsigned prediction errors were significantly coded in a right superior frontal cluster, spanning the superior and middle frontal gyri (peak: 27 8 54, cluster size: 145 voxels, T=8.43, Z=7.68, p_FWE_<.001), a left superior frontal cluster spanning the superior and middle frontal gyri (peak: −26 0 54, cluster size: 32 voxels, T=7.66, Z=7.09, p_FWE_<.001), and a dorsal anterior cingulate cortex cluster (peak: −7 12 50, cluster size: 6 voxels, T=5.32, Z=5.1 p_FWE_=.002 Fig. 3A). We also found a right superior occipital cluster (peak: 24 -74 34, cluster size 21 voxels, T=7.24, Z=6.75, p_FWE_<.001) and two parietal clusters: (peak: 34 -48 38, cluster size: 18 voxels, T=5.69, Z=5.43, p_FWE_<.001) (peak: 20 -60 26, cluster size: 4 voxels, T=5.24, Z=5.03, p_FWE_=.003). We used the left and right SFC and dACC clusters as ROIs with which to take forward our analysis of dopaminergic effects on precision weighting in our primary analyses. Secondary analyses examined these effects in occipital and parietal regions.

**Figure 3:**
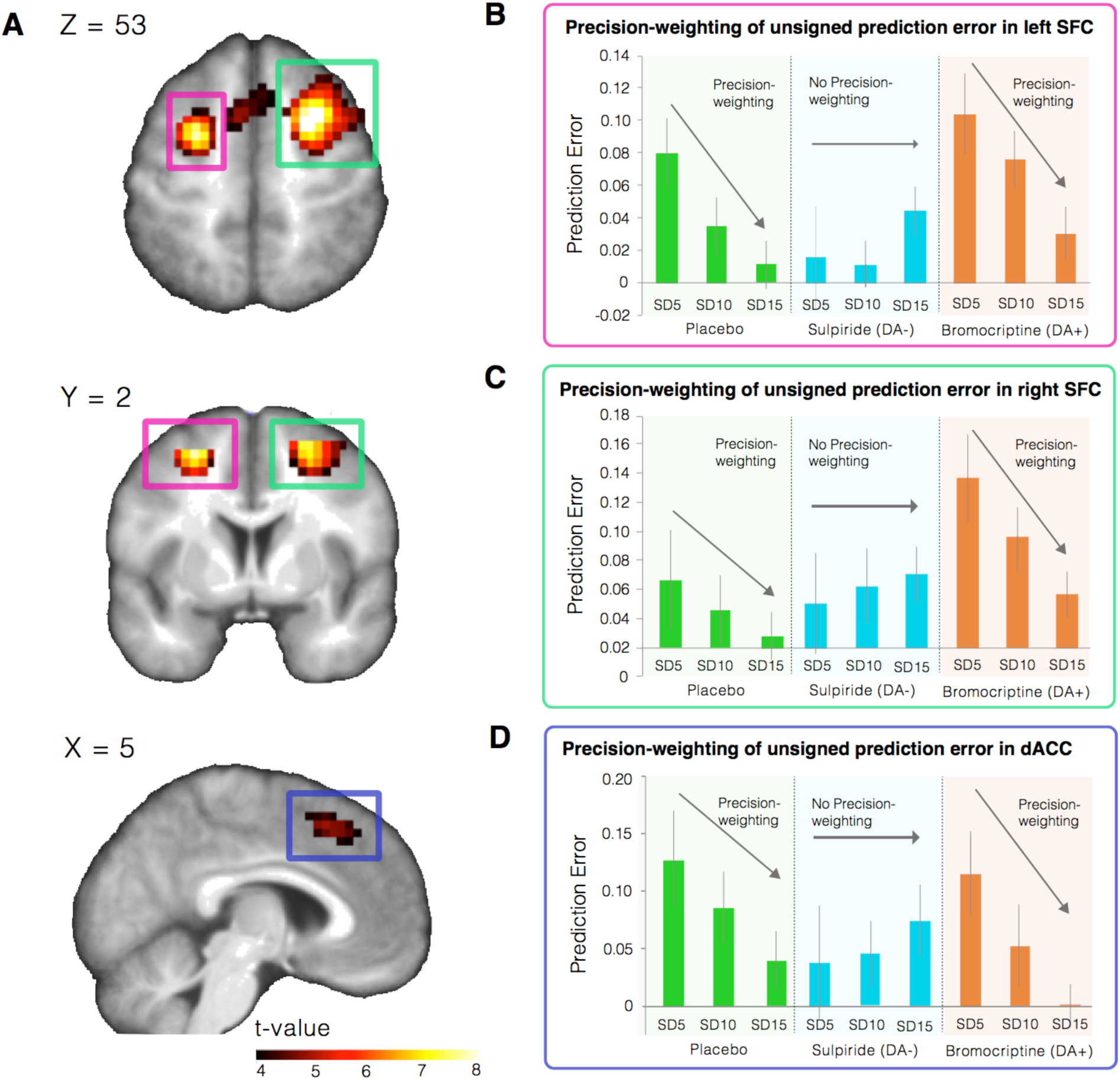
A. Unsigned prediction errors were coded in bilateral superior frontal cortex and dorsal anterior cingulate cortex. The left side of the brain is the left side of the image. B-D. When exploring these regions further, we find that unsigned prediction errors are coded in a precision-weighted fashion as indicated by the strong unsigned prediction error signal in the SD5 condition which declines over the SD10 & SD15 condition in the placebo and bromocriptine group. Importantly, sulpiride perturbed precision-weighting significantly in the left SFC. Error bars represent standard error of the mean.

### Precision-weighting of unsigned prediction errors is mediated by dopamine in the SFC/dACC

In order to test for an effect of outcome precision (i.e., the SD of reward distributions) and pharmacological perturbations on precision-weighting of unsigned prediction errors, we extracted the parameter estimates (betas) of the unsigned prediction error parametric modulators from the left and right superior frontal cortex (SFC) and dACC cluster that showed a main effect of unsigned prediction error coding at whole brain corrected *p*FWE<.01. We used a two-factor mixed model ANOVA with medication group as the between-subjects variable and SD condition as the within-subjects variable, using a linear contrast across SD for the main effect of SD and interaction. In the left SFC cluster, there was a significant interaction across SD conditions and medication group, suggesting indicating that medication group had a significant effect on precision-weighting of unsigned prediction errors (F{2,56}=4.025 *p*=.023; Fig. 3B). We then explored whether the effect was driven by differing effects of bromocriptine or sulpiride compared to placebo. In the analysis of differential effects of sulpiride and placebo, we found a significant interaction between medication group and SD condition (F{2,37} = 5.44, *p*=.025), which indicates that sulpiride dampens precision-weighting of unsigned prediction errors. Comparing the placebo and bromocriptine group, there was a significant effect of SD (F{1,37} = 14.93, *p*<.001), but no significant effect of group (F{1,37} = 2.781, *p*=.104) or interaction between medication group and SD (F{2,36} = 0.02, *p*=.894). This indicates that SFC unsigned prediction error signals are precision-weighted, but unaffected by bromocriptine. In the right SFC we did not find a significant interaction (F{2,56}=1.70, *p*=.193; Fig. 3C), and therefore did not perform any further post-hoc tests. However, the brain response pattern is largely the same as in the left SFC (see Fig. 3).

In the dACC we found a “trend-level”, marginally significant interaction (F{2,56}=2.81, *p*=.069; Fig. 3D). Post-hoc tests between the placebo and sulpiride group revealed a trend-level interaction (F{1,38}=3.043, *p*=.089). Testing the placebo and bromocriptine group revealed a significant effect of SD (F{1,37}=9.32, *p*=.004), but no effect of group (F{1,37}=1.17, *p*=.29) and interaction (F{1,37}=.172, *p*=.68). A similar pattern was thus found in the dACC as in the left SFC (see Fig. 3).

### Performance correlates with the precision-weighting of unsigned prediction errors

As neural precision-weighting is thought to facilitate task performance, we computed Spearman correlations between the degree of precision-weighting (quantified as the average parameter estimates (betas) of the unsigned prediction error in the SD5 condition minus the SD15 condition) and performance (quantified as the mean performance error (abs(actual mean – predicted mean)) on the last three trials in each session. Thus a lower performance (error) error equals higher accuracy (performance)). We found that precision-weighting in both the left superior frontal cortex (Rho=-.43, *p*=.001; Fig. 4A) and the right superior frontal cortex (Rho=-.37, *p*=.004; Fig. 4B) was significantly related to performance, and precision-weighting in the dACC was trend-level related to performance (Rho=-.24, p=.066), such that increased precision-weighting resulted in more accurate predictions (smaller difference between prediction and actual mean). In a general linear model controlling for medication group this effect remained significant (F{57}=7.53, *p*=.008).

**Figure 4:**
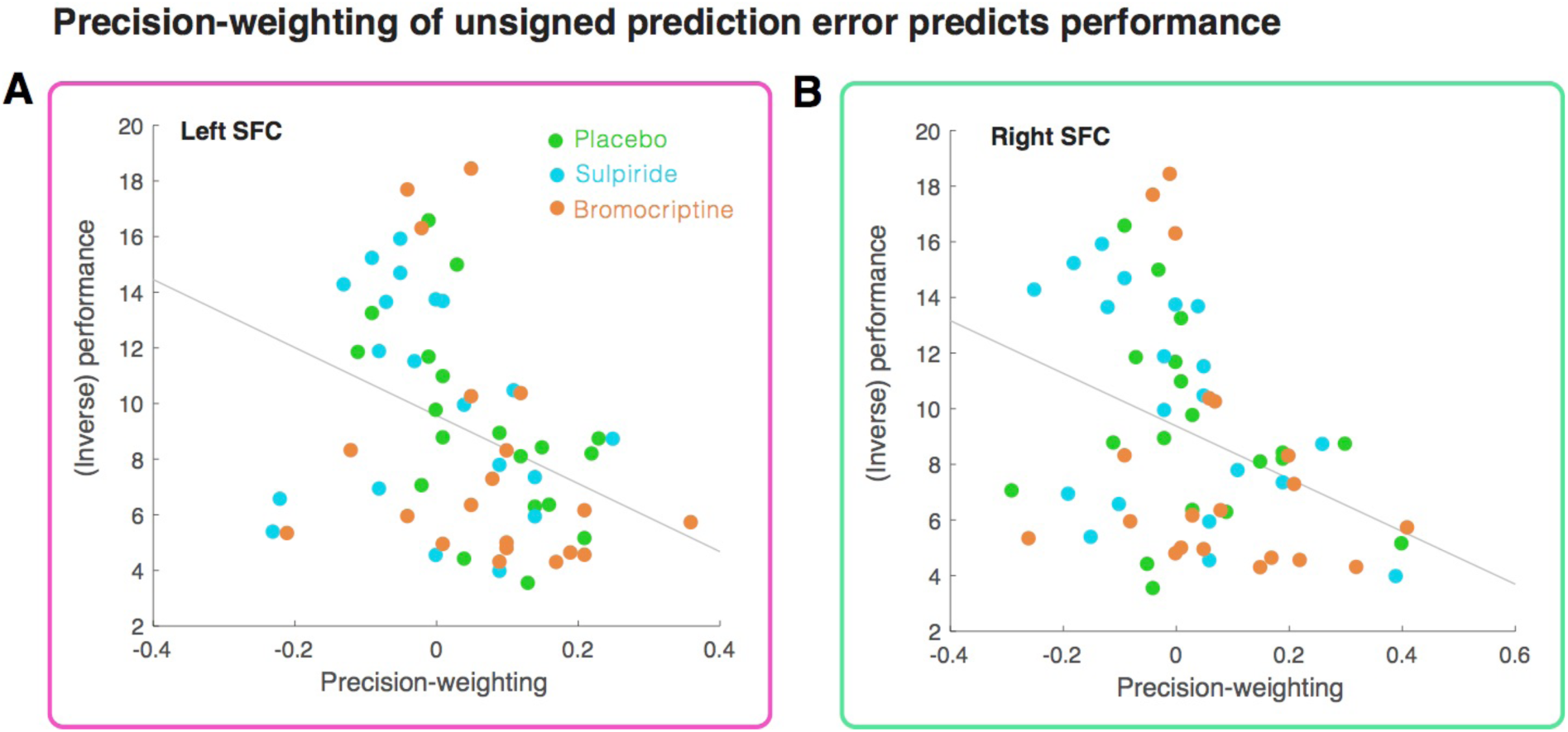
Precision-weighting of unsigned prediction errors in the A. left SFC and B. right SFC correlates with performance (i.e. difference between mean of the reward distribution and predicted mean) on the task.

### Secondary outcome variable analyses

In addition to the analyses we conducted in our primary regions of interest, i.e. SFC and dACC, we also test for an interaction between SD conditions and medication in the remainder of clusters that coded unsigned prediction errors. There were no significant SD by medication interactions: superior occipital cortex cluster, (F{2,56}=1.60, *p*=.21), larger parietal cluster (F{2,56}=.82, *p*=.45), smaller parietal cluster (F{2,56}=1.53, *p*=.23).

## Discussion

We first confirmed that unsigned prediction errors play a role in learning using a regression analysis. We subsequently confirm using computational modelling on the behavioural level that a separately estimated precision-weighted signed and unsigned prediction errors provided the best description of the behavioural data, thus suggesting precision-weighted signed and unsigned prediction errors in the brain (the former confirmed previously by fMRI in humans (Diederen et al 2016), including in a prior analysis of these data (Diederen et al., 2017). In the present study, we tested the prediction of brain representation of unsigned prediction errors. We found evidence supporting this prediction, by showing that unsigned prediction errors are coded relative to the SD in an associative learning paradigm (i.e. precision-weighted) in the bilateral superior frontal cortex and dorsal anterior cingulate cortex (SFC/dACC). Importantly, this mechanism was significantly diminished in the sulpiride (D2 receptor antagonism) group in comparison to the other groups in the right SFC, with marginal evidence of a medication effect in the dACC. This result suggests that the dopamine D2 receptor plays a key role in the mechanisms underlying precision-weighting of unsigned prediction errors. We furthermore found that a decrease in precision-weighting of unsigned prediction error was significantly correlated to performance on the task, where an increase in precision-weighting resulted in more accurate predictions of upcoming rewards. These results confirm the hypothesis that there exist cortical unsigned prediction error signals, which influence performance and are precision weighted by dopamine.

Although we argue that these results are in line with influential predictive coding theories of cognition, the reinforcement learning models used in this paper are not hierarchical. This is because an important element in hierarchical Bayesian models, like the hierarchical Gaussian filter model (Mathys et al., 2011), lies in the ability to learn about higher level features of the reward environment that were absent in this task. In our task, participants were explicitly informed that the distance between the green bars in the cues indicated the relative standard deviation of the reward distribution (low, medium high), although they were uninformed with regards to the exact size of the standard deviation (SD5, SD10 & SD15). Furthermore the mean of the distribution remained fixed within each session, i.e. there were no changes in the mean and/or the volatility of changes. The experimental manipulation introduced by the task design eliminates the need to estimate latent variables of interest and examine their fit with MRI data. Rather, simple reinforcement-learning models and classical neuroimaging contrasts suffice to model this task. An advantage of making the precision of reward information explicit is that we can test directly the degree of precision-weighting, which would otherwise be implicit, as in the hierarchical Gaussian filter, where precision-weighting is a consequence of the precision of – or confidence in –higher-level priors. Although the models are not hierarchical in this paper, we believe these results are in line with influential formulations of predictive coding and the proposed role of dopamine therein (Refs – Friston 2009, Friston et al 2014). In our view, traditional experimental psychology task design and latent variable “model-based” fMRI are complementary ways of investigating brain function.

Although sulpiride diminished precision-weighting on the neural level, there were only mild deficits in performance induced by sulpiride, and our behavioural modelling suggested that on average, the sulpiride group still utilised precision-weighting in their decision making. The potential reasons for this include that our analysis on the behavioural level may not have been sufficiently sensitive to demonstrate sulpiride-induced impairments (Murray et al., 2010). Alternatively, a secondary mechanism in the brain might compensate for the lack of superior frontal cortex precision-weighting of prediction error in the sulpiride condition, potentially at the cost of performance in more taxing circumstances (Murray et al., 2010).

It could be objected that a difference in brain encoding of unsigned prediction errors might be driven by different sizes of prediction error across SD conditions. For example, it could be objected that in the SD15 condition unsigned prediction errors will be higher than in the SD5 condition, as it is a more noisy environment. Critically, our findings are the reverse of what would be expected under this objection. That is, whilst the objection posits higher brain prediction error associated activity in the SD15 condition because prediction errors are higher here than in the SD5 condition, our data indicates the opposite effect: that prediction errors in the SD15 condition are in fact attenuated compared to the SD5 condition.

The coding of unsigned prediction errors in the superior and middle frontal gyri and dACC is in line with earlier findings by Hayden (et al., 2011) who found unsigned prediction errors in the dACC of monkeys, and with prior fMRI studies in humans (Fletcher et al 2001, Turner et al 2004, Fouragnan et al., 2017; Fouragnan et al 2018; Metereau et al., 2013; Ide et al., 2013). However, the finding that cortical unsigned prediction error signals are precision-weighted in humans is a novel finding to the best of our knowledge. The fact that such a precision weighted signal exists in the dACC is notable given that, aside from the motor cortex, dopaminergic innervation of cortex is greatest in ACC (Lewis et al., 1987; Berger et al., 1991; Paus, 2001), compatible with the hypothesis that precision weighting is influenced here by dopaminergic input.

The observation that dopamine plays a role in the precision-weighting of unsigned prediction errors extends previous work showing the precision-weighting of signed (reward) prediction errors (Diederen et al., 2016 & 2017), and fits well with contemporary computational neuroscience models which suggest that the brain is a hierarchically organized system which aims to minimize prediction error. According to these accounts, modulatory neurotransmitters like dopamine are hypothesised to play a key role in controlling the precision of prediction errors (Bastos et al., 2012; Adams et al., 2013; Friston et al., 2014a,b). Our finding that the precision of the prediction error signal is represented in superior and middle frontal gyri and anterior cingulate cortex under placebo, and that the degree of precision-weighting of the signal is disrupted by sulpiride, provides initial evidence in accordance with such models. We did not find evidence that bromocriptine modulated precision-weighting of cortical prediction error signals. This is analogous to the results of the previous analysis of these data focussed on signed prediction error signals, especially in subcortical regions. Diederen and colleagues (2017) found that there were effects of sulpiride on precision of the prediction error signal in the midbrain and striatum, but found no corresponding effect of bromocriptine. Choosing the optimal dose of medication to use in pharmacological fMRI studies is often a challenge, where tolerability has to be balanced against efficacy. Furthermore, bromocriptine has been shown to have non-linear dosage effects. For example a study investigating motor cortex neuroplasticity under different dosages of bromocriptine (2.5mg, 10mg and 20 mg) found a non-linear dose relationship (Fresnoza et al., 2014), and effects of dopaminergic drug intervention may depend on baseline dopamine levels (Cools et al., 2011). Thus, it is possible that the dose of bromocriptine used in this study (2.5mg) may have been too low to detect robust effects on brain signals, and thus its lack of significant effect on precision weighting in this experiment should not be over-interpreted.

If the precision-weighting of prediction errors is important in learning, we can expect aberrant learning to occur when prediction errors are not scaled optimally to the precision of information. This mechanism could be of importance to psychosis, which is characterized by delusional beliefs and hallucinatory perception (Fletcher & Frith et al., 2009). Previous work showed aberrant prediction error coding in people with psychosis (Murray et al., 2008; Corlett et al., 2007a,b). As psychosis has consistently been associated with dopamine dysfunction (Howes et al., 2009), it is conceivable that the precision-weighting process is impaired in psychosis. Indeed, it has been suggested that dopamine dysregulation causes psychosis due to affecting the brains capacity to precision-weight prediction error (Adams et al., 2013). That is, if unreliable prediction errors were given excessive weight, they could have an exaggerated influence on driving changes in the brain’s model of the world, thereby contributing to the formation of abnormal beliefs. As such, future studies should aim to explore the relationship between precision-weighting of prediction error and psychosis.

In conclusion, we found evidence of precision-weighted unsigned prediction errors in the superior frontal and dorsal anterior cingulate cortices. Furthermore, we found that the precision-weighting of prediction errors was modulated by the dopaminergic antagonist sulpiride, and we found that the degree of precision-weighting in this area was correlated to performance on the task, providing evidence for the first time that dopamine plays a role in precision-weighting of unsigned prediction error signals during learning.

## Acknowledgements

We would like to thank Wolfram Schultz for his important role in designing the experiment and acquiring the data analysed here, and to the staff of the Clinical Research Facility and Wolfson Brain Imaging Centre, Cambridge Biomedical Research Centre for assistance in data collection.

Author roles: J.H.: Conceptualization, methodology, formal analysis, writing (original draft preparation, review and editing); P.C.F.: Conceptualization, project administration, funding acquisition, writing (review and editing). T.S. and H.Z. investigation, project administration, writing (review and editing). K.M.J.D. Conceptualization, methodology, project administration, implementation, formal analysis, supervision, writing (original draft preparation, review and editing) G.K.M., conceptualization, project administration, methodology, supervision, writing (original draft preparation, review and editing)

Conflicts of Interest: P.C.F. has received payments in the past for ad hoc consultancy services to GlaxoSmithKline All other authors declare no competing interests.

Funding: This work was supported by the Neuroscience in Psychiatry Network, a strategic award from the Wellcome Trust to the University of Cambridge and University College London (095844/Z/11/Z), Wellcome Trust (093270) Bernard Wolfe Health Neuroscience Fund (P.C.F. and H.Z)., and the Niels Stensen Foundation (K.M.J.D).

## Methods

### Demographics

|  | N | Mean Age (SD) | Male (Female) |
| --- | --- | --- | --- |
| **Medication study** | 59 | 24.19 (4.46) | 31 (28) |
| *Placebo* | 20 | 23.9 (4.76) | 9 (11) |
| *Sulpiride* | 20 | 24.8 (4.51) | 12 (8) |
| *Bromocriptine* | 19 | 23.7 (4.3) | 10 (9) |

### Participants

We recruited 63 participants, of which 4 were excluded due to feeling unwell during scanning and/or a failure to complete the paradigm, or side-effects of the medication (see Diederen et al., 2017 for a detailed description). All participants were recruited via the distribution of flyers in Cambridge and advertisements on Gumtree. Drug screenings were all negative. After receiving detailed information about the study, all participants gave written informed consent.

### Pharmacological dopaminergic challenge

Participants received a single dose of the D2-antagonist sulpiride (600mg), the dopamine agonist Bromocriptine (2.5 mg), or placebo, in a double-blind fashion. Domperidone was added to prevent potential nausea, as it would be an indicative factor of having taken a drug, and could thus influence the blinding procedure, as well as making people feel unwell (see Diederen et al., 2017 for a detailed description of the dopaminergic manipulation).

### Task design

Durin functional (f)MRI data acquisition, participants predicted the magnitude of upcoming rewards. Optimal performance on this task required the participant to estimate the mean of the distribution from which the rewards were drawn. The task consisted of three sessions of 10 minutes each. The rewards were drawn from six different pseudo-Gaussian distributions that differed with respect to their standard deviation (SD) and expected value (i.e. mean of the distribution; EV). See Table 1 for an overview of the SD’s and EV’s. Distributions were counterbalanced to ensure that the two conditions within each session differed with respect to the EV and SD. The conditions were presented in short blocks, each including 4–6 trials. Each distribution consisted of 21 trials, which resulted in a total number of trials of 42 per block. The participants were informed beforehand that each distribution (of which two per session) had a different level of precision, which could be one of three levels: low, medium or high precision, corresponding to SD5, 10 & 15, although the exact SD’s were not revealed to the participants. Each trial started with a fixation cross that was presented between 2100 and 4200ms. The fixation cross was followed by a cue that was presented for 500ms. This cue informed the participant from which of two distributions (high, medium or low) in that block the upcoming reward was being drawn. After the cue was presented the participants were required to predict the magnitude of the upcoming reward. They had 3500ms to make this prediction. Making the prediction involved scrolling a mouse ball with their fingers up and down, and clicking the left mouse button to state their prediction. This resulted in moving a bar up and down the screen on a scale that ranged from 0 to 100. The starting point on the bar was randomized throughout the experiment, to ensure that scrolling distance did not correlate with participants’ predictions. The participants were instructed to minimize the prediction error, i.e. the difference between the expected and the actual obtained reward. After the prediction, another fixation cross was presented for duration between 2100 and 5250ms. Thereafter, the obtained reward was presented in addition to the prediction and the reward prediction error for 1000ms (see Fig. 5 for an example trial).

**Table 1:** Overview AIC for each reinforcement learning model per medication group. The lowest AIC for each group is printed in bold.

|  | Free parameters | AIC overall | Placebo | Sulpride | Bromocriptine |
| --- | --- | --- | --- | --- | --- |
| RW | 1 | 894.7 | 824.2 | 935.9 | 924.0 |
| RW scaled | 1 | 903.2 | 829.0 | 947.3 | 933.4 |
| PH not scaled | 2 | 884.2 | 808.2 | 931.1 | 913.4 |
| PH both scaled | 2 | 883.6 | 807.3 | 930.1 | 913.4 |
| PH-estimated SD | 3 | 882.7 | 806.1 | 928.9 | 914.1 |
| PH-estimated SD separate for signed and unsigned PE | 4 | **879.4** | **803.1** | **926.3** | **910.4** |

**Table 2:** Participants

|  | N Sessions | N Trials | SDs | EVs | N participants |
| --- | --- | --- | --- | --- | --- |
| **Dopamine Study** | 3 | 62 | 5, 10 & 15 | 35 & 65 | 60 (20 per group) |

**Figure 5:**
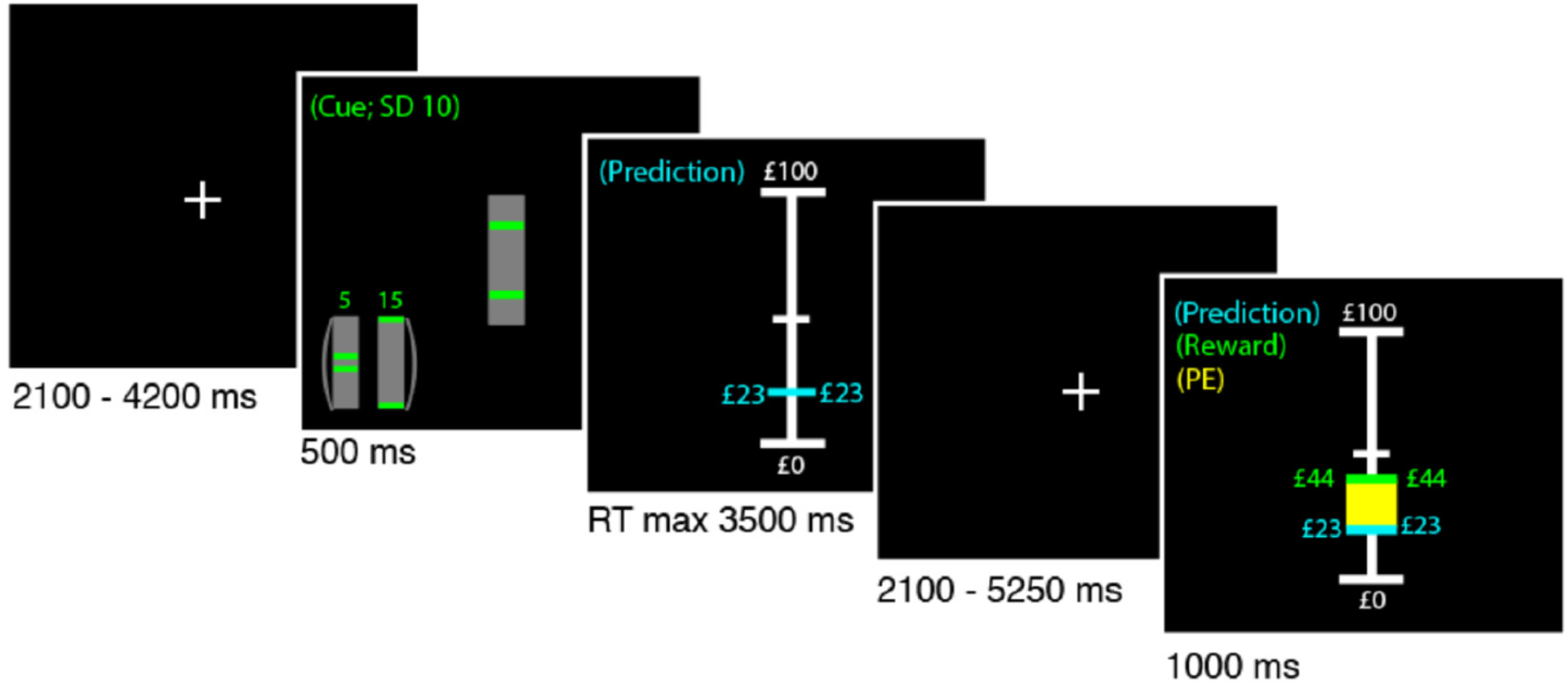
Example of a trial. The participants were instructed to learn the mean of a reward distribution. First a fixation cross was presented after which the participants were informed about the standard deviation of the reward distribution. Subsequently the participants were asked to make a prediction regarding the upcoming reward, which was presented to the participant in combination with the prediction error (in yellow) after an anticipation period.

### Pay-off

If the participants would always receive the amount of money drawn by the computer, this might have reduced their motivation to reveal their true prediction. Therefore, the participants were rewarded according to their accuracy (i.e., proximity to the mean of the reward distribution) on 20% of the trials in order to keep them motivated to predict as well as possible.

### Behavioural analyses

Behavioural data was analysed in Matlab. In order to test the performance of the participants, we analysed whether the predictions of the participants approached the mean of the distribution (or expected value (EV)), toward the end of the experiment (final 3 predictions in each SD condition). We calculated the absolute difference between the EV and the prediction in these final 3 trials and averaged them. We reasoned that if participants’ prediction were close to the EV of the reward distribution from which rewards were drawn, the participants were learning. We tested this by using a mixed-2-factor-ANOVA with a 3-level within subject factor (SD5, SD10 & SD15) and a 3-level between subject factor (medication), and tested for main effect of SD and interaction between SD conditions and medication using a mixed-model ANOVA in SPSS (version 21). Specifically we used a linear contrast (termed “linear polynomial contrast” in SPSS) for the main effect of SD and interaction. In addition, we analysed differences in scrolling distance to exclude the possibility that drug effects influenced willingness to do the task, resulting in a decrease in scrolling distance. We also analysed differences in reaction times and missed trials for the same reason.

### Linear regression model

In order to explore the contribution of signed and unsigned prediction error to learning we used a linear regression model with belief-updating as dependent variable and both unsigned and signed prediction error as independent variables. Belief updating here is defined as the change in value estimate from t_n_ to t_n+1_. That is, when the participant expects a value of 20 on t_1_ and expect as value of 30 on t_2_ there was a belief update of 10, which is expected to be explained by a prediction error experienced at t1. Since signed prediction errors, but not unsigned prediction errors inform the participant about the direction of learning, we expect the former but not the latter to have a main effect on belief-updating. Furthermore, since formal learning models like the Pearce-Hall model (Pearce & Hall, 1980), suggest that unsigned prediction errors increase learning, we expect an interaction between unsigned and signed prediction error on belief-updating. In order to visualise the effect of signed and unsigned prediction errors on belief-updating, we will create bins on the basis of signed prediction errors and plot belief-updating on the y-axis. If unsigned prediction errors interact with signed prediction error in predicting belief-updating we should expect to see a logit or sigmoid relationship between signed prediction error and belief updating.

### Behavioural modelling

To investigate learning, we fitted several reinforcement learning models to participants’ prediction sequences. Each model used a common updating rule in which predictions on trial *n* depended on the prediction error and the learning rate on trial *n-1:*

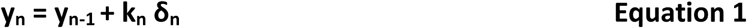

Here, y_n_ is the prediction made on trial *n*, *k_n_* refers to the learning rate and *δ_n_* denotes the size of the prediction error. This is a standard reinforcement learning model, and allows to estimate to which degree prediction errors are being used by the participant by estimating the learning rate parameter (Sutton & Barto et al., 1998).

The first model consisted of a Rescorla-Wagner (RW) reinforcement learning model with a fixed learning rate. A fixed learning rate prescribes that each prediction error is weighted equally during learning (e.g., independent of whether a prediction error occurred at the start, or the end of a session):

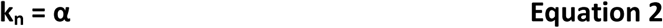

The second model consisted of a Pearce-Hall (PH) model with a trial-wise, dynamic, learning rate, which prescribes higher weighting of prediction errors (i.e., more learning) at the start of a task session compared to later trials (Pearce et al., 1980). In uncertain environments, it is more optimal to decrease the weighting of prediction errors as learning progresses (once participants become more certain of their predictions) as prediction errors will continue to occur as a result of the imposed uncertainty. The PH learning rate decays as trial progress, as a function of the previous learning rate and the experienced prediction error, which allows for fast updating of predictions at the beginning of each task session:

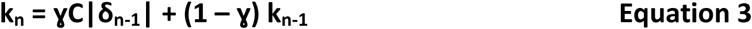
in which |δ| is the absolute prediction error, C is an arbitrary scaling coefficient and γ is the learning rate decay.

We additionally explored whether scaling prediction error to the reliability of the environment (i.e., precision –weighting) benefitted learning by comparing models that scaled the prediction error term and models that did not. We both explored models that simply precision-weighted the prediction error by a constant term ω (equation 4 & 5), and models that estimated the degree υ of precision-weighting for each individual (equation 6). We estimated both models that had a single estimated u for both signed and unsigned prediction errors and a model that had a separate variable for signed and unsigned prediction errors. In contrast to earlier studies (Diederen et al., 2015, 2016 & 2017) we used a more simplified model that utilised a linear precision-weighted model to ease interpretation.

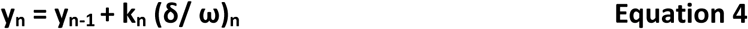

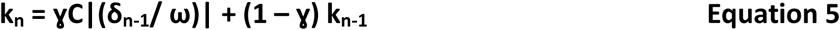

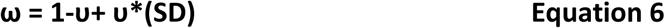

For the Pearce-Hall model we explored the scaling of the signed and unsigned prediction error separately, as well as combined.

We used the Akaike Information Criterion (AIC) to estimate the model evidence in favour of each of the two models. The model with the lowest AIC was used for further analyses.

### fMRI pre-processing steps

All pre-processing steps were performed using SPM8 (available at http://www.fil.ion.ucl.ac.uk; Wellcome Department of Cognitive Neurology, London, England) in combination with the Donders matlab (dmb) toolbox for combining data from different echo acquisition times. Data was acquired at 4 different echo times (TE’s) of 12, 27,91, 43,82 and 59,73 ms. Corrections for slice timing was not applied considering the relatively fast repetition times (TR) of 2100ms. Using SPM8 the functional images were realigned first, after which the four different echoes were summed and averaged in order to minimize signal intensity inhomogeneity due to differences in T2* relaxation times across the brain (Poser et al., 2006). The functional images were subsequently coregistered to the T1-weighted anatomical image in native-space by first coregistering one averaged functional image, after which the coregistration parameters of this registration were applied to the all the functional images in order to registered to native space. Unified segmentation was used in order to achieve the normalization of the functional and anatomical images to MNI space (Ashburner et al., 2005). Unified segmentation is a single iterative model that combines segmentation, bias correction and normalization in order to achieve optimal results. The segmentation step was omitted for one of the participants; since it gave erroneous segmentation parameters resulting in distortions in the normalized anatomical T1 scan. The segmentation step was later conducted using a different template, namely the East-Asian template, which resulted in a correct segmentation. Spatial smoothing was performed using 8mm Gaussian kernels. The time series in each session were high-pas filtered at 128 Hz.

### 1^st^-level analysis

fMRI analyses were conducted using SPM8. We used a single statistical linear regression model for our main analyses. We modelled for each trial the onset of the cue as an event (i.e. a delta function of zero duration), the onset of the prediction event (i.e., when participant could start making their prediction) as a single epoch lasting until they indicated their prediction. Each event was convolved by the standard canonical haemodynamic response function in SPM8. Importantly, we used parametric modulation to identify neural correlates of unsigned prediction error responses by specifying for all outcome events the reward prediction errors. The reward events were separately modelled for the different SD conditions in order to test for differences in precision-weighting of prediction errors as evidences by different sizes of slopes for the coding of unsigned prediction errors under different levels of certainty. Contrast were created on the 1^st^-level which combined the two EV conditions for each SD in order to leave a single contrast for each SD conditions which could be taken to the 2^nd^-level (i.e., group-level).

### 2^nd^-level analysis

For the analysis of the main effect of unsigned prediction errors we first created 1^st^-level contrast in which we combined the 2 EV’s for each separate SD condition, resulting in 3 contrasts corresponding to the 3 different SD conditions (SD5, SD10 & SD15). We modelled these 3 SD contrasts for each pharmacological condition separately specifying the effects to be dependent, and having equal variance. We subsequently tested for a main effect of prediction error by doing a positive 2^nd^-level contrast on the parametric modulators for the unsigned prediction error.

To test for an effect of SD on the coding of unsigned prediction error and the effect of pharmacological group, we extracted the parameter estimates (beta’s) for all SD conditions for each medication group separately from the left and right superior frontal cortex clusters and dACC cluster surviving FWE<.01 coding unsigned prediction errors; for each cluster we used the mean values per cluster per participants in a mixed-2-factor-ANOVA with a 3-level within subject factor (SD5, SD10 & SD15) and a 3-level between subject factor (medication), and tested for main effect of SD and interaction between SD conditions and medication using a mixed-model ANOVA in SPSS (version 21). Specifically we used a linear contrast (termed “linear polynomial contrast” by SPSS) across SD to examine the main effect of SD and interaction between group and SD. When a significant interaction was found we explored this further to see which group was driving the interaction.

